# From Flies to Humans: A Conserved Role of CEBPZ, NOC2L, and NOC3L in rRNA Processing and Tumorigenesis

**DOI:** 10.1101/2025.01.11.632529

**Authors:** Guglielmo Rambaldelli, Valeria Manara, Andrea Vutera Cuda, Giovanni Bertalot, Marianna Penzo, Paola Bellosta

## Abstract

NOC1, NOC2, and NOC3 are evolutionarily conserved nucleolar proteins that play an essential role in the maturation and processing of ribosomal RNA (rRNA). NOC1, in *Drosophila* is necessary to sustain rRNA processing, whereas its depletion leads to impaired polysome formation, reduced protein synthesis, and induces apoptosis. In this study, we demonstrated that the RNA-regulatory functions of NOC1 are conserved in vertebrates, where the reduction of the CEBPZ homolog of NOC1 leads to the accumulation of unprocessed 45S pre-rRNA, a reduction in protein synthesis, and inhibition of cell growth. Gene Ontology and bioinformatic analyses of CEBPZ, NOC2L, and NOC3L in tumors highlight a significant correlation between their expression and processes that regulate rRNA processing and ribosomal maturation. Moreover, comparative analysis of TCGA datasets from tumor databases revealed that CEBPZ, NOC2L, and NOC3L exhibit contrasting expression patterns across tumor types. This context-dependent behavior suggests that overexpression of these proteins may promote tumor growth, whereas reduced expression could exert tumor-suppressive effects, underscoring their complex and unexpected regulatory roles in cancer.

**Summary Statement:** NOC1, NOC2, and NOC3 are conserved nucleolar proteins essential for rRNA processing. Their reduction impairs rRNA maturation, decreases protein synthesis, and induces cell death. CEBPZ, NOC2L, and NOC3L show context-dependent expression in tumors, suggesting dual roles in cancer progression and suppression.

## Introduction

Ribosome biogenesis is a highly regulated process that relies on specific nucleolar proteins to facilitate the processing of ribosomal RNA (rRNA), which is essential for proper ribosome maturation. Disruptions in this pathway can result in aberrant ribogenesis and contribute to proliferative disorders, including tumor initiation (Dorner et al., 2023; Hwang and Denicourt, 2024; Penzo et al., 2019). Nucleolar Complex Proteins 1, 2, and 3 (NOC1, NOC2, and NOC3) are evolutionary conserved nucleolar proteins essential for growth in *S. cerevisiae* and *Arabidopsis* (Edskes, Ohtake and Wickner, 1998; Milkereit et al., 2001) (Li et al., 2009). Their function was first characterized in yeast for their ability to form functional NOC1/2 and NOC2/3 heterodimers, which are necessary for the processing and transport of the rRNA during the maturation of the 60S ribosomal subunit (Milkereit et al., 2001). In addition, NOC proteins are critical for maintaining nucleolar integrity and facilitating the incorporation of ribosomal proteins into ribosomes (Dorner et al., 2023; Hurt, Iwasa and Beckmann, 2024; Sanghai et al., 2023; Vanden Broeck and Klinge, 2023), ensuring proper ribosome biogenesis. NOC proteins are also present in *Drosophila*, where we have demonstrated their essential role in controlling organ and animal growth (Destefanis et al., 2022). Furthermore, we showed that NOC1 controls rRNA processing, and its reduction results in rRNA accumulation, disrupts polysome assembly, and compromises protein synthesis (Destefanis et al., 2022). Additionally, mass spectrometry analysis of the NOC1 interactome identified NOC2 and NOC3 among the interacting proteins, suggesting that NOC1 may also form heterodimers with NOC2 and NOC3 in *Drosophila,* as observed in yeast (Manara et al., 2023).

In vertebrates, CEBPZ, NOC2L, or NOC3L are essential genes, and in mice, their respective knockout are embryonic lethal (Friedel et al., 2007). Additionally, data from DepMap, a platform that reports genetic dependencies in cancer (Fong et al., 2024; Meyers et al., 2017; Tsherniak et al., 2017), indicates that they are all essential for tumor growth (Ciani et al., 2022).

CEBPZ (ID10153) belongs to the C/EBP (CCAAT protein) family of transcription factors (Lum et al., 1990). Known for its role as a DNA-binding transcriptional activator of heat shock protein 70, CEBPZ’s function in vertebrates remains largely unexplored. In tumors, CEBPZ has been proposed as a marker for colorectal and gastric cancers, while point and missense mutations have been identified in patients with acute myeloid leukemia (AML) (Herold et al., 2014), where CEBPZ works as a co-factor to maintain m6A modifications in leukemia cells (Barbieri et al., 2017).

NOC2L, or NIR (Inactivator of histone acetylases), is a nucleolar histone acetyltransferase inhibitor that represses the transcription of genes controlling the cell cycle (Li et al., 2021). Conditional knockout mice exhibit increased p53 levels, apoptosis, and bone marrow defects (Ma et al., 2014). Interestingly, silencing of NOC2L/NIR results in pre-rRNA accumulation and increases p53 levels (Wu et al., 2012). Depletion of NOC2L in LOVO tumor cells inhibits their growth in xenograft mice (Li et al., 2021), supporting a role in tumor progression. Moreover, elevated NIR levels in colorectal cancer serve as a marker for malignancy in patients with poor disease outcomes (Li et al., 2021).

NOC3L or Fad24 (Factor for adipocyte differentiation-24), located in the nucleolus, is required for the adipocyte differentiation of NIH-3T3-L1 cells (Tominaga et al., 2004). While its overexpression is linked to colorectal and gastric carcinomas, its reduction suppresses the in vitro growth of gastric cancer cells (Yan et al., 2020), suggesting a similar behavior to that reported for NOC2L in LOVO cells (Li et al., 2021). Notably, in zebrafish, NOC3L/Fad42 reduction significantly affects the development of hematopoietic cells (Walters et al., 2009), mirroring our unpublished observation on CEBPZ reduction in zebrafish, suggesting a role in the differentiation of hematopoietic cells. NOC3L has also been shown to compromise DNA replication and induce apoptosis in HeLa cells, suggesting an additional role in regulating transcription and pre-initiation complex (PIC) formation (Cheung et al., 2019).

This study uncovers a conserved and critical function for the NOC proteins in ribosomal biogenesis. In *Drosophila and* yeast, NOC1, NOC2, and NOC3 are indispensable for properly processing rRNA. We propose that CEBPZ/NOC1, NOC2L, and NOC3L collaboratively regulate rRNA maturation in humans, and that disruption of anyone leads to defective rRNA processing, impaired ribosome assembly, and cell death. GO analysis revealed that CEBPZ, NOC2L, and NOC3L are associated with RNA processing and rRNA maturation. Notably, co-expression analysis of CEBPZ and NOC2L showed a significant enrichment in genes that participate in R-loop metabolism. These support a recent study identifying CEBPZ as part of the protein complexes associated with R-loop resolution in human stem cells (Wu et al., 2021). R-loops are structures of RNA-DNA hybrids normally formed during transcription, also present in the nucleolus due to the high rate of transcription of rRNAs (Feng and Manley, 2022; Petermann, Lan and Zou, 2022; Wells, White and Stirling, 2019).

Expression profiling across diverse tumor types revealed distinct patterns for CEBPZ, NOC2L, and NOC3L, elevated in several cancers, while markedly reduced in acute myeloid leukemia (AML) and kidney chromophobe carcinoma (KICH). This distinctive and intriguing pattern correlates with poor prognosis in specific cancer types and suggests a context-dependent, cell–type–specific role for these genes in tumor progression. Notably, this variation may be related to their potential involvement in regulating R-loops, pointing to a broader role in maintaining genomic stability beyond ribosome biogenesis.

## Results

### 1. NOC proteins control rRNA processing in flies and human cells

Ubiquitous reduction of NOC1, NOC2, or NOC3 in *Drosophila* impairs animal development, with NOC1 reduction affecting protein synthesis and inducing apoptosis (Destefanis et al., 2022). Here, we show that this effect is common for all three NOCs, as reduction of either NOC1, NOC2, or NOC3 affects the correct maturation of rRNAs, as shown by the accumulation of ITS1 and ITS2 pre-rRNA forms (Figure 1A-B) and by reduction of the mature 18S and 28S ribosomal RNAs (Figure 1C-D). In addition, we also found up-regulation of p53 at transcriptional levels (Figure 1E), supporting the increase in apoptosis observed previously, as a response to stress (Destefanis et al., 2022).

**Figure 1.**
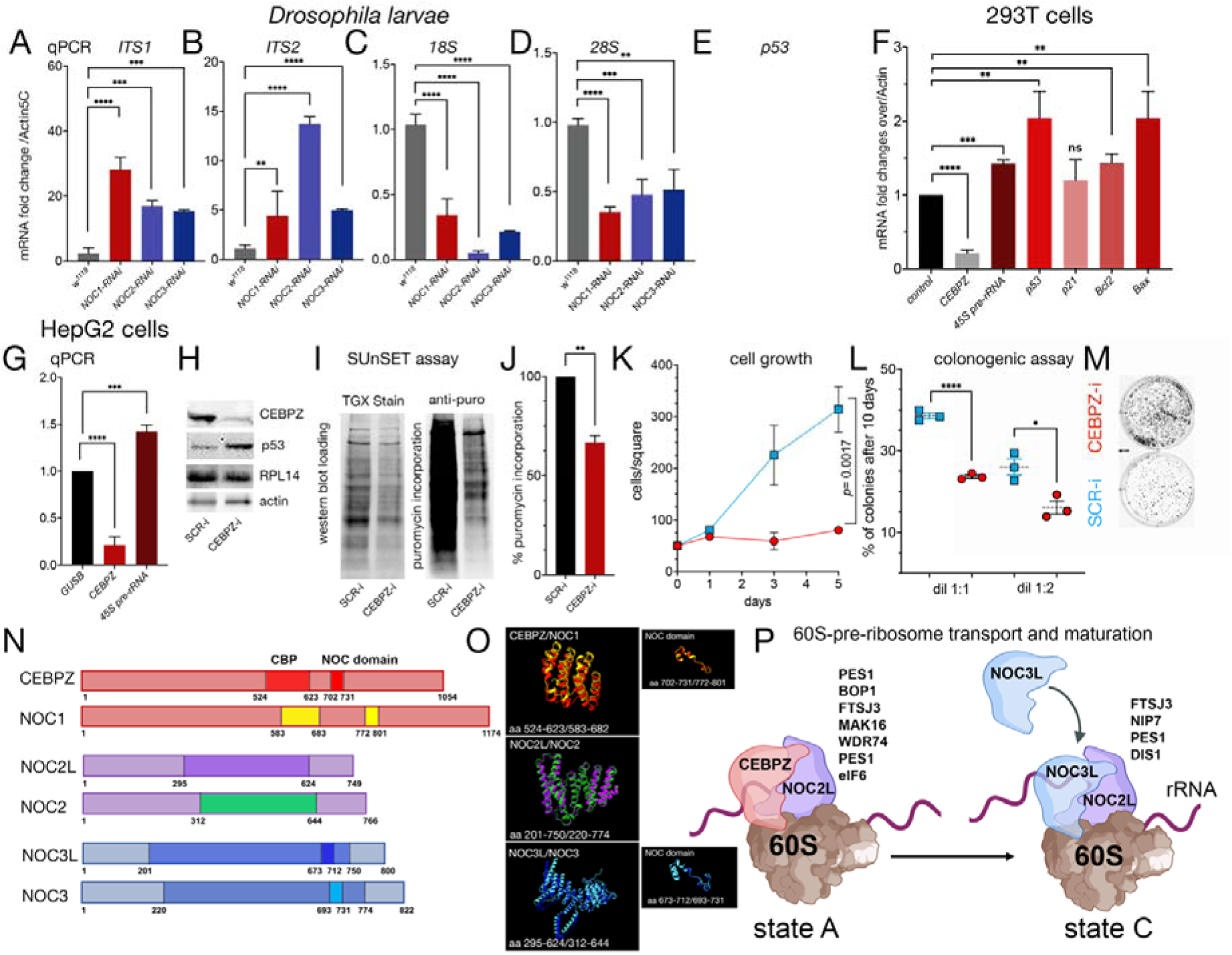
**(A-F) Reduction of Nucleolar Complex proteins (NOCs) in Drosophila leads to rRNA accumulation, impaired maturation of 18S and 28S rRNA, and elevated p53 levels**. (A-E) qRT-PCRs from *Drosophila* whole larvae with ubiquitous reduced *NOC1, NOC2,* or *NOC3* showing an accumulation of pre-rRNAs, analyzed using the ITS1 (internal transcribed spacers) 1 and 2 (A), and a reduction of 18S and 28S rRNA expression (C-D) and of *p53 mRNA* (E). The level of *NOC1, NOC2,* or *NOC3-mRNAs* is calculated over control *w^1118^* animals; data are expressed as fold increase relative to control *Actin5C;* RNA interference efficiency is shown in Supplementary Figure 1. **(F-M) Reducing CEBPZ in human tumor cells results in rRNA accumulation, impaired protein synthesis, and decreased cell growth.** (F-G) qRT-PCR analysis of HEK 293FT cells (F) and HepG2 cells (G) following siRNA-mediated silencing of CEBPZ, showing increased levels of 45S pre-rRNA and p53 mRNA. Expression levels are presented as fold change relative to control, normalized to β-glucuronidase (GUSB). Data from (A-G) are presented as mean ± s.d. for at least three independent experiments \*\**p<*0.01; \*\*\**p<*0.001; \*\*\*\**p<*0.0001 (using Student’s t-test for the analysis). (H) Western blot from HepG2 lysates of cells transfected with a scramble siRNA (SCR-i) or for siCEBPZ (CEBPZ-i) showing the level of CEBPZ, p53, RPL14 used as an unrelated protein, and actin as a control loading. (I) SUnSET assay; western blot from cells treated with 1 µg/ml of puromycin shows the relative changes in protein synthesis using anti-puromycin antibodies in cells treated with siSCR, or siCEBPZ, total protein loading is shown using stain-free technologies (TGX Stain-Free Fastcast). (J) Quantification of the relative change in the rate of puromycin incorporation between SCR-i and CEBPZ-i data was analyzed using two independent experiments. (K-L) (K) Cell growth expressed at day 1,3 and 5, and (L) percentage of colonies at 10 days after treatment with siCEBPZ and siCTL data are expressed from two separate experiments where cells were originally plated at two different concentrations \**p<*0.05; \*\*\*\**p<*0.0001 (using Student’s t-test for the analysis). (M) Photos of colonies at 10 days of treatment. (N-O) Schematic representation of *Drosophila* and humans NOC1/CEBPZ, NOC2/NOC2L, and NOC3/NOC3L proteins. CEBPZ contains a CBP (CCAAT binding domain in dark RED) with 32% amino acid-sequence identity with *Drosophila* NOC1, and a conserved NOC domain (in red and yellow). This domain is also present in NOC3L and NOC3 (highlighted in blue and azure). NOC2 shares an overall 36% of amino acid identity between *Drosophila* and human NOC2L, with the highest homology (47%) represented with a box (in purple and green); see sequence homologies in Supplementary File 2. **(O) The predicted structural homology of the conserved regions between CEBPZ/NOC1, NOC2L/NOC2, and NOC3L/NOC3** was obtained by a simulation using AlphaFold and ChimeraX analysis. **(P) Schematic representation of the CEBPZ-NOC2L and NOC2L-NOC3L heterodimers at states A and C of the 60S ribosomal maturation**. The graph also includes proteins identified through DepMap analysis (Table 1) due to their significant correlation with the expression of the corresponding heterodimers. Their role in the ribosomal complex is illustrated according to the maturation state (A and C) of the 60S ribosome subunit as described in the literature (Vanden Broeck and Klinge, 2023; Vanden Broeck and Klinge, 2024). Figure 1P was realized using Biorender, license EY27N2G66G.

**Table 1.**
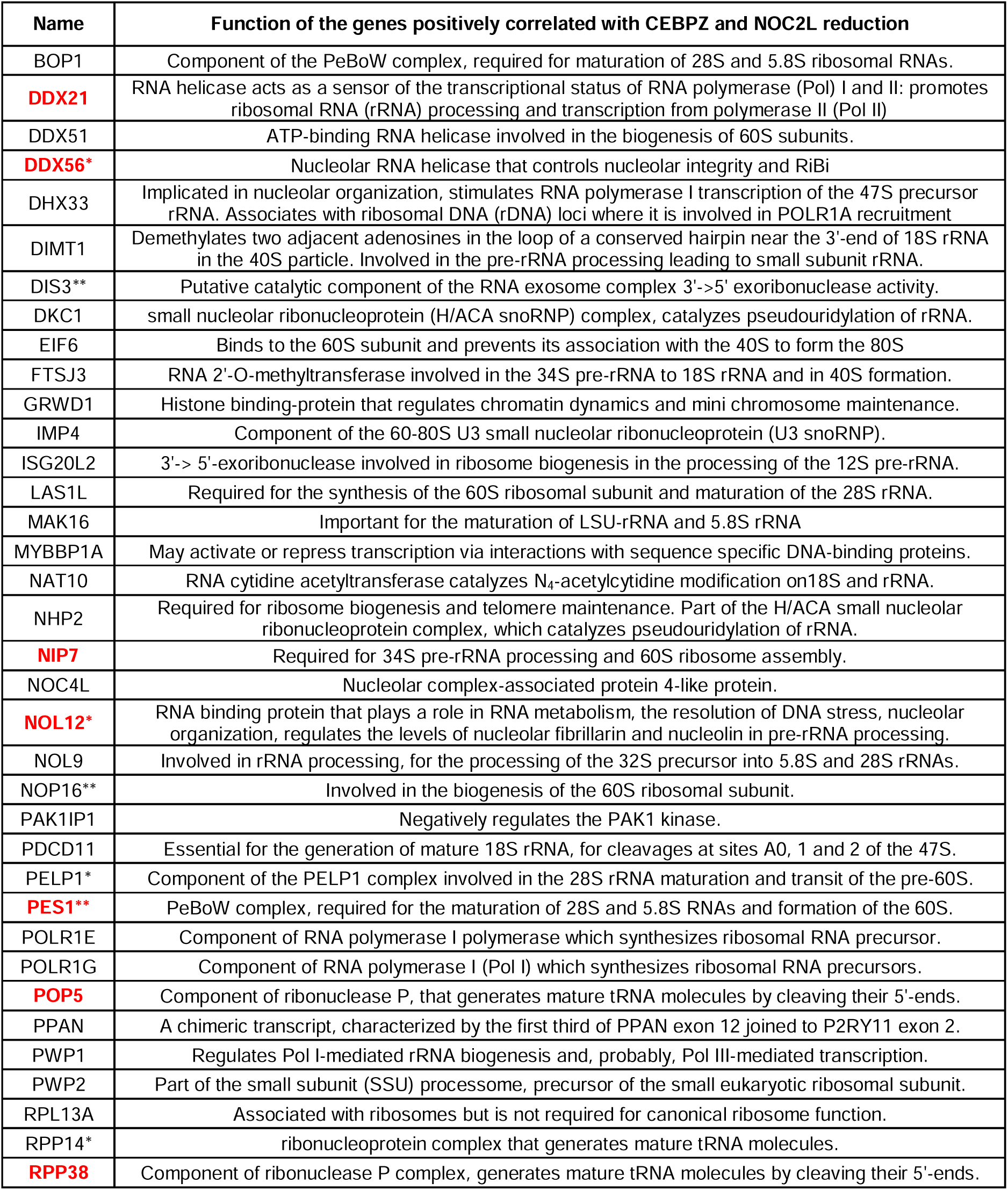
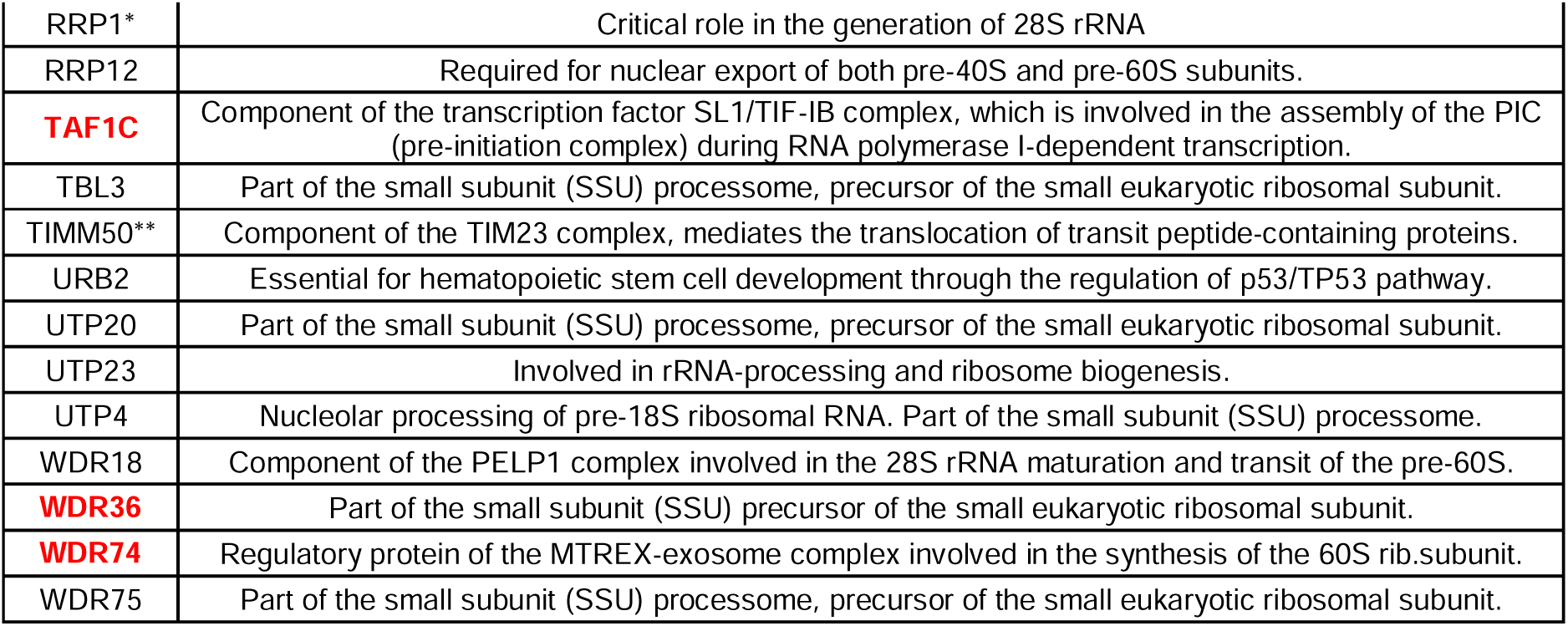
Genes positively correlated with CEBPZ and NOC2L upon their reduction in tumor cells and their functional annotation from the UniProt database (UniProt, 2024). * Positively correlating when CEBPZ and NOC3 are reduced; ** positively correlate when all three genes are reduced. The whole list of genes is available in Supplementary File 3. In red are genes associated with R-loop formation and metabolism.

This mechanism appears conserved in humans, as reducing the human homolog CEBPZ in HEK293T tumor cell line induces the accumulation of *45S pre-RNA* and increases the levels of *p53 mRNA,* and its target *Bcl2* and *Bax* (Figure 1F). Similar results were observed in the HepG2, liver carcinoma cell line, where CEBPZ reduction also leads to increased levels of *45S pre-RNA* (Figure 1G) and elevated p53 protein level (Figure 1H). SUnSET assay shows a significant reduction in puromycin incorporation in cells silenced for CEBPZ, indicating decreased protein synthesis (Figure 1I-J). This reduction correlates with impaired cell growth, as shown by the clonogenic assay (Figure K-M). These data recapitulate the observations in *Drosophila* and support the hypothesis that CEBPZ plays an evolutionarily conserved role in regulating rRNA maturation.

Human NOC proteins share 32% amino acid identity with their *Drosophila* homologs (Figure 1N). Analysis to predict their 3D structures using Alphafold (Jumper et al., 2021; Varadi et al., 2022) imported into ChimeraX (Goddard et al., 2018; Meng et al., 2023) revealed the presence of highly similar structures in the proteins of the two species, with 37, 47, and 40 % of identity for CEBPZ, NOC2L, and NOC3L, respectively (Figure O). Both CEBPZ and NOC1 contain a CAAT-binding protein (CBP) and the NOC domains, the last also present in NOC3L and NOC3 (Figure 1N-O).

Based on these conserved structural similarities, we hypothesized that CEBPZ may be part of the complex in which heterodimers with NOC2L participate in the transport and cleavage of rRNA and maturation of the 60S ribosomal subunit, and a model has been created to explain these passages (Figure 1P) (Hurt, Iwasa and Beckmann, 2024). During these maturation events, heterodimers CEBPZ-NOC2L are formed and in the state A with proteins including PES1, BOP1, FTSJ3, and WDR1, which we found in our GO analysis for CEBPZ-NOC2L co-expressing genes, significantly correlating with their expression (Table 1 and in Figure 1P). In further maturation steps, CEBPZ detaches from NOC2L, allowing NOC3L to bind and heterodimerize with NOC2L in a complex necessary at state C of the 60S ribosomal maturation (Hurt, Iwasa and Beckmann, 2024). In addition to FTSJ3 and PES1, this complex contains NIP7 and DIS3, which were found to be coregulated with NOC2L-NOC3L in our expression analysis (Table 1 and in Figure 1P).

### 2. Gene Ontology analysis links CEBPZ, NOC2L, and NOC3L depletion to rRNA processing and ribosome biogenesis in tumor cells

To explore the functional significance of CEBPZ, NOC2L, and NOC3L in cancer, we analyzed data from the Cancer Dependency Portal (https://depmap.org/portal/), which utilizes CRISPR screening to identify genes that are critical for tumor growth.

Starting from the CRISPR knockdown data from DepMap, we extracted and performed our analysis on the top hundred co-expressed genes for each of the three genes of interest (Supplementary File 3). We then selected the shared co-expressed genes between the three. We performed GO enrichment analysis to examine coregulated pathways and identify the functional implications of our genes and the possible contributions of individual heterodimers and the assembled complex.

The GO enrichment analysis of the shared genes reveals a network of interconnected biological processes, predominantly in ribosome biogenesis, RNA processing, and rRNA maturation, which are enriched across all three data sets (Figure 2A-C). Network visualization (Figure 2D) from these data highlights the functional clustering of genes involved in RNA metabolism and ribosomal assembly. The network illustrates the functional interdependence of these processes, implying that perturbations in CEBPZ, NOC2L, or NOC3L can significantly impact ribosomal function and rRNA processing. From this analysis, it is plausible that decreasing the expression levels of CEBPZ or NOC2L disrupts the formation of CEBPZ-NOC2L heterodimers, leading to defects in rRNA accumulation and affecting the maturation of the ribosomal subunits. Additionally, although to a lesser extent, we found that reduction of the NOC2L-NOC3L interaction affects RNA processing and ribosome biogenesis, reinforcing our hypothesis that both pairs play a role in regulating rRNA maturation, and impairment of either heterodimer disrupts rRNA maturation, leading to its accumulation.

**Figure 2.**
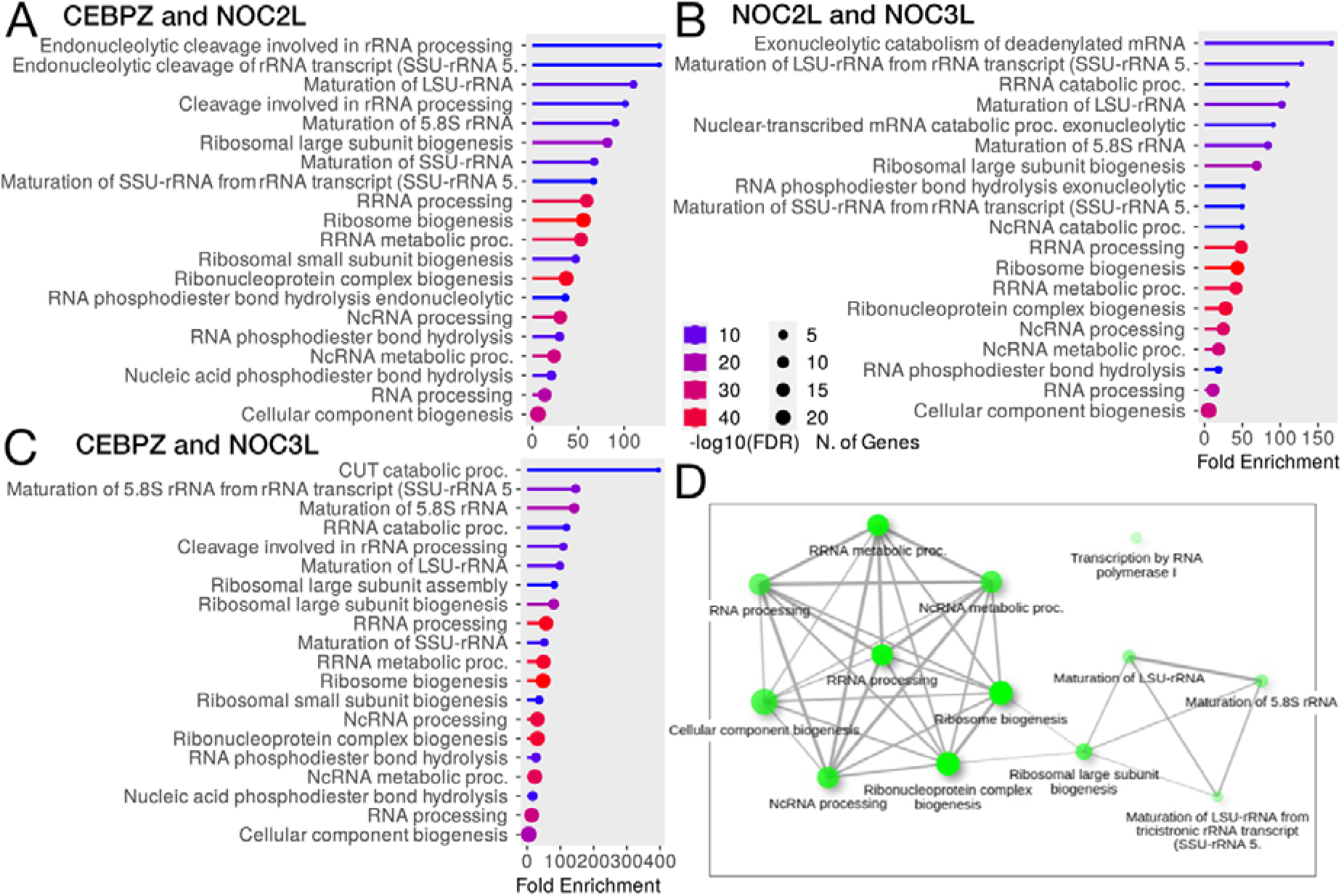
GO enrichment of the shared co-regulated genes between CEBPZ, NOC2L, and NOC3L. (A-C) GO enrichment (Biological Processes) of the shared co-regulated genes between the two indicated target genes. The results are ordered based on the Fold Enrichment; the circle size represents the number of genes in the pathway, the color of the bars the -log10(FRD), and the length of the bar the enrichment. (D) GO network representation. Brighter nodes are more significantly enriched gene sets. Bigger nodes represent larger gene sets. Thicker edges represent a higher number of genes shared between the two sets.

In Table 1, we show the list of genes that positively correlate with CEBPZ and NOC2L reduction, and we highlight in red the genes associated with R-loop formation of metabolism (22% of the total), excluding CEBPZ.

### 3. CEBPZ, NOC2L and NOC3L tumor expression analysis

Dysregulation of rRNA processing is a hallmark of cancers. We analyzed the expression across diverse tumor types based on the hypothesis that CEBPZ, NOC2L, and NOC3L form regulatory heterodimers that modulate this process. Specifically, we examined their expression levels using TCGA data from the UALCAN cancer portal (Chandrashekar et al., 2017; Chandrashekar et al., 2022) (Figure 3A-C). This analysis revealed that while all three genes were significantly upregulated (*P*<0.05) in most tumors (Figure 3D, in red), their expression was significantly reduced (*P*<0.05) in acute myeloid leukemia (LAML), and Kidney Chromophobe carcinoma (KICH) (Figure 3D, in green). In contrast, in KIRC and KIRP renal carcinomas, while the expression of CEBPZ was low, that of NOC2L and NOC3L increased (Figure 3D). These data provide evidence of different controls for the expression of these genes in different cell-tissue contexts. Indeed, as these genes may be upregulated in many solid tumors, to support increased ribosome production and protein synthesis for rapid cellular proliferation, their selective downregulation in hematopoiesis and kidney tumors KICH may suggest a tissue-specific or cell-specific regulation. Moreover, the fact that NOC2L and NOC3L are upregulated in KIRC/KIRP may indicate that they may exert a dual function, perhaps in transcriptional regulation or cell cycle control, beyond rRNA processing.

**Figure 3.**
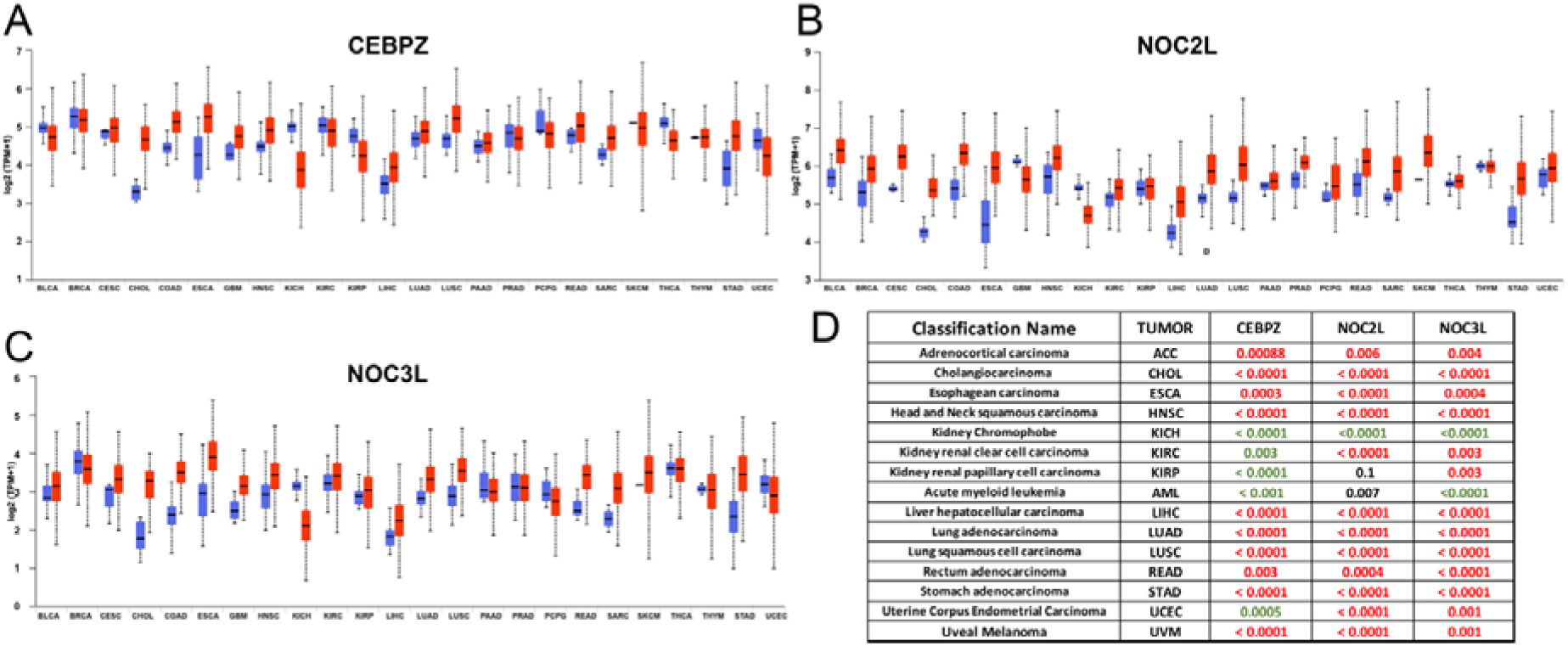
Gene expression analysis (TCGA). (A-C) Expression levels of *CEBPZ* (A), *NOC2L* (B), and *NOC3L* (C) in tumor (red) versus normal (blue) tissues across various cancer types. (D) Summary panel highlighting cancer types where the expression of at least two of the three genes is significantly altered (*P* < 0.05), as reported in the UALCAN portal. Red indicates overexpression, while green indicates downregulation in tumor tissues relative to normal control.

We then examined CEBPZ, NOC2L, and NOC3L expression patterns across various stages of tumor development using TGCA datasets from the UALCAN portal. This analysis revealed a significant increase in their expression in liver hepatocellular carcinomas (LICH) and lung adenocarcinoma (LUAD) (Figure 4A-B). The observed upregulation correlated with the progression of tumor stages. The expression levels of LIHC in stage 4 were comparable to those in controls, possibly because of the reduced sample size for this stage. Similar upregulation was found for rectum adenocarcinoma (READ) and stomach adenocarcinoma (STAD) (not shown), suggesting that CEBPZ, NOC2L, and NOC3L overexpression may contribute to the aggressiveness of these carcinomas, probably by sustaining the high rate of rRNA processing and ribosome activity in these tumors. In contrast, in kidney chromophobe carcinomas, CEBPZ, NOC2L, and NOC3L expression levels were consistently and significantly reduced (Figure 4C), and variable in other kidney cancers (Figure 4D-E), supporting the conclusion of our previous analysis.

**Figure 4.**
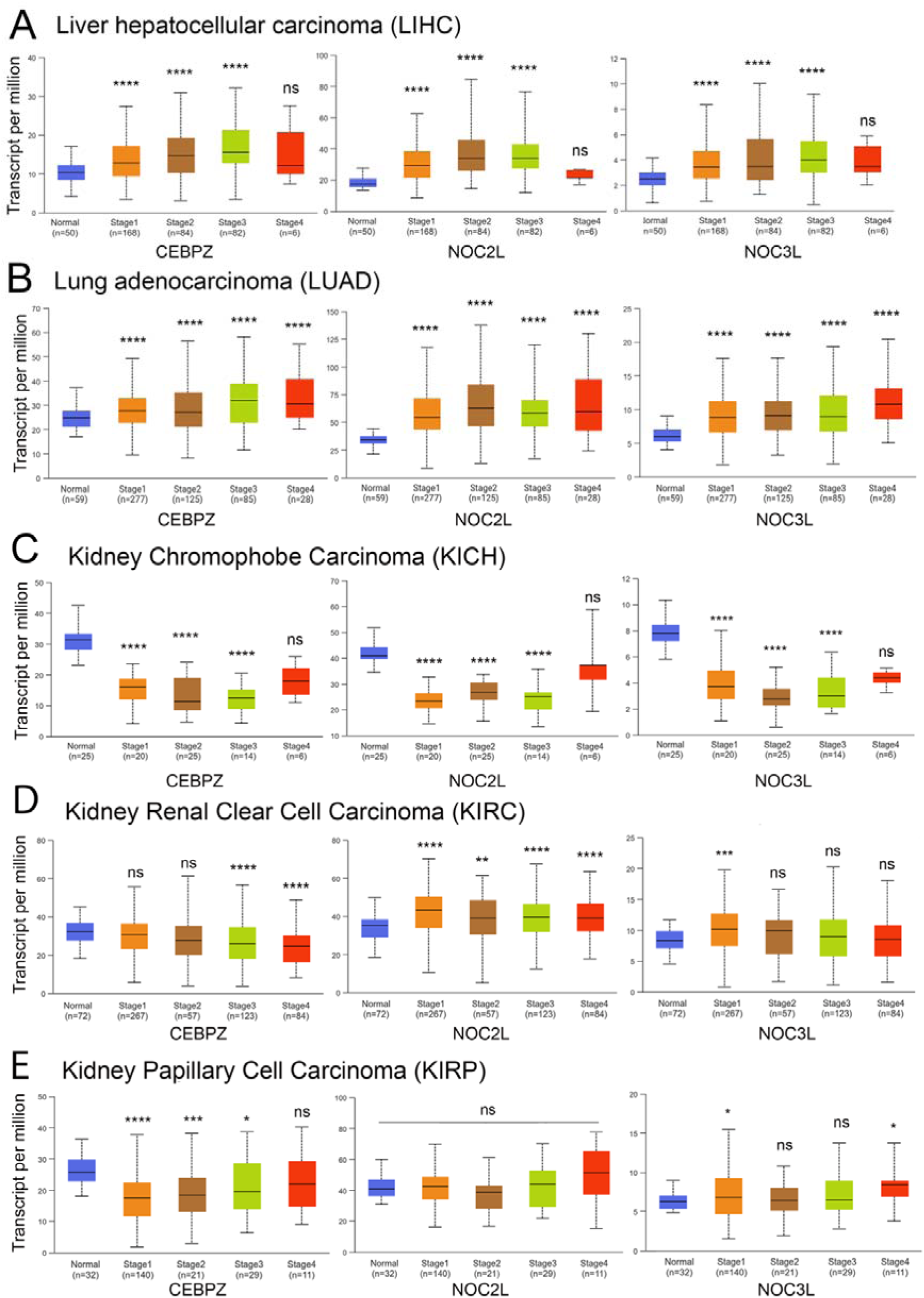
Graphs illustrating expression levels across various cancer types during tumor progression. Data from UALCAN highlights the correlation between gene expression and disease progression, demonstrating significant changes in the expression of CEBPZ, NOC2L, and NOC3L with advancing disease stages. (A) Liver hepatocellular carcinoma (HCC) and (B) Lung adenocarcinoma (LUAD), (C) Kidney Chromophobe Carcinoma (KICH). (D) Kidney Renal Clear Cell Carcinoma (KIRC). (E) Kidney Papillary Cell Carcinoma (KIRP). The statistical analysis was performed by comparing the expression levels of the indicated genes in tumors at various stages to their expression levels in normal control tissue; statistical significance is expressed by the asterisks * *p*< 0.05, ** *p*< 0.01, *** *p*< 0.001, **** *p*< 0.0001, ns: not significant.

To deeply investigate the co-expression patterns of CEBPZ, NOC2L, and NOC3L within kidney tumors, we analyzed the TCGA datasets publicly accessible through FireBrowse (Cerami et al., 2012). A Pearson correlation coefficient was calculated between the expression of the three genes (CEBPZ, NOC2L, and NOC3L) in the three kidney carcinoma subtypes (KICH, KIRC, and KIRP). This analysis showed that CEBPZ and NOC3L exhibited the highest direct correlation in all three tumors (Table 2). Meanwhile, a weak negative correlation existed between CEBPZ and the other two genes in KIRC and KIRP. This suggests their potential distinct regulation across various kidney cancers.

**Table 2.**
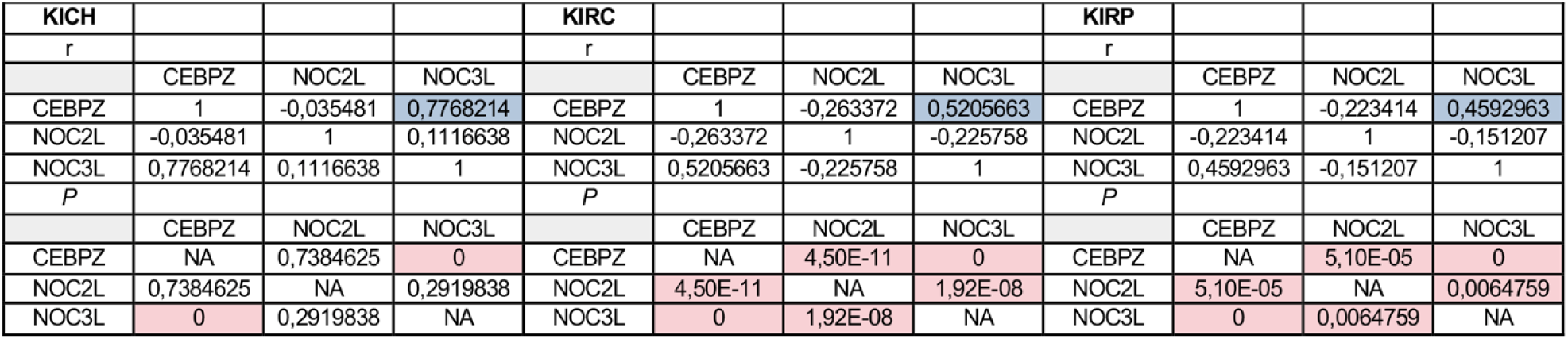
A matrix highlighting the significant correlation between CEBPZ, NOC2L, and NOC3L expression in KICH, KIRC, and KIRP kidney carcinomas. It shows that for each tumor type, the r (correlation coefficient) and the *P* value, highlighted in blue, are significant, with a correlation higher than -/+ 0.4. A higher absolute number means a stronger correlation between the samples’ expression levels. (e. g. in KICH, when CEBPZ is low, NOC3L is also low). The lower part of the table shows the *P*-value for each correlation, highlighting the significant ones (*P*<0.05).

### 3.2 Impact of CEBPZ, NOC2L, and NOC3L expression on patient survival

Finally, we analyzed the correlation between the expression of CEBPZ, NOC2L, and NOC3L and patient survival to explore their potential impact on disease outcomes across tumor types. Figure 5 displays a heatmap of hazard ratios for these genes, providing a visual summary of how their expression is related to survival in various tumors.

**Figure 5.**
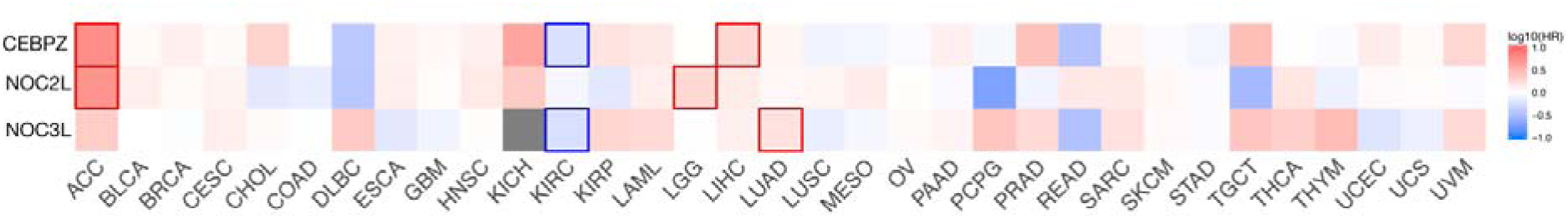
Hazard ratio (HR) heat map for CEBPZ (ID: ENSG00000115816.13), NOC2L (ID: ENSG00000188976.10), and NOC3L (ID: ENSG00000173145.11) in the tumors listed below. The median is selected as a threshold for separating high-expression and low-expression cohorts. The bounding boxes depict the statistically significant results (*P* < 0.05). Colors from white to red indicate an HR above 1, suggesting that high gene expression is associated with a worse prognosis. Colors from white to blue indicate an HR below 1, suggesting that high gene expression is associated with a better prognosis.

These data suggest that the expression of CEBPZ and NOC2L in adenoid cystic carcinoma (ACC) may be associated with poorer tumor prognosis. This means that as the expression levels of these genes increase, the severity or aggressiveness of cancer may also increase, potentially leading to worse patient outcomes. A similar trend was observed in brain lower-grade glioma (LGG), liver hepatocellular carcinoma (LIHC), and lung adenocarcinoma (LUAD), with different degrees of significance (Figure 5). In contrast, there was an inverse correlation between CEBPZ and NOC3L expression and patient outcomes in kidney chromophobe carcinoma. Additionally, CEBPZ, NOC2L, and NOC3L exhibited distinct expression patterns during the progression of kidney carcinoma, with a marked decrease observed in KICH (Figure 4C). These findings reinforce our observation that impairing RNA processing and metabolism in ribosome biogenesis, through downregulation of CEBPZ, NOC2L, and NOC3L, alters key cellular pathways, potentially contributing to tumor progression in a cell-type-specific manner, particularly in kidney-derived cancers such as KICH.

## Discussion

This study reveals a conserved and novel role for the human nucleolar proteins CEBPZ, NOC2L, and NOC3L in ribosome biogenesis. Our initial analysis of the NOC1 protein interactome in flies identified NOC2 and NOC3 proteins as components of the NOC1-nucleolar complex (Manara et al., 2023). These findings sustain studies in yeast, where NOCs were shown to form NOC1-NOC2 and NOC2-NOC3 heterodimers necessary for rRNA processing and 60S ribosome maturation (Dorner et al., 2023; Milkereit et al., 2001).

Based on these observations, we hypothesized that in humans, CEBPZ, NOC2L, and NOC3L also function as heterodimers to control rRNA maturation as illustrated in the model in Figure 1P. However, although NOC1 has been previously characterized in yeast as a factor primarily involved in the maturation of the 60S ribosomal subunit, our data reveal that its depletion also impairs the processing of 18S rRNA, a component of the 40S subunit. The simultaneous reduced 18S and 28S rRNA maturation upon NOC1 reduction indicates a more global effect on ribosomal RNA processing and points to a central, conserved function of NOC1 in coordinating the assembly and maturation of ribosomal components. Knockdown of CEBPZ leads to pre-rRNA accumulation and activation of p53, consistent with a canonical nucleolar stress response (Lindstrom, Bartek and Maya-Mendoza, 2022). This upregulation of p53 was first observed in *Drosophila* and linked to a DNA damage response upon NOC1 reduction (Pederzolli et al., 2025). Moreover, we find p53 upregulated when each of the NOC genes was reduced (Figure 1E), an event that may suggest that reduction of each of the three NOCs may induce p53 upon ribosomal stress to activate the surveillance pathways. We propose a similar mechanism for the vertebrate homologs, CEBPZ, NOC2L, and NOC3L, where the role of p53 as a downstream effector also emphasizes the potential tumor-suppressive consequences of NOC dysfunction in some tumors. The dual roles of NOC proteins in supporting tumor growth (via ribosome biogenesis) and potentially triggering p53-mediated arrest when disrupted highlight their context-dependent functions. This complexity may account for the variable expression patterns observed across different cancer types and their mixed prognostic significance.

The association between NOCs’ aberrant expression and tumor progression suggests that dysregulation of NOC protein expression could contribute to oncogenesis (Figure 4). TGCA and GO analyses show that CEBPZ, NOC2L, and NOC3L expression levels correlate with genes relevant to rRNA maturation and ribosome processes. Thus, their overexpression may enhance ribosome biogenesis to sustain tumor progression (Table 1).

Differential expression of CEBPZ, NOC2L, and NOC3L across cancer types with varying prognostic implications implies tissue-specific or context-dependent roles. Indeed, their elevated expression in certain cancers correlates with poor prognosis (Figure 5). Conversely, reduced expression in other contexts may reflect a tumor-suppressive role or vulnerability in specific oncogenic settings. Paradoxically, in kidney carcinomas and acute myeloid leukemia, the reduced levels of NOC proteins may impair ribosomal function, triggering cell-type–specific apoptosis or may activate senescence programs (Nishimura et al., 2015; You and Wu, 2025), where this selective pressure could favor the emergence of aggressive clones through the accumulation of mutations that bypass these fail-safe mechanisms. A similar trend in the altered expression of other ribosome-biogenesis factors, such as dyskerin and fibrillarin, has been observed in tumorigenesis (Penzo et al., 2017; Zhang et al., 2024). Fibrillarin upregulation correlates with poor prognosis in breast cancer and AML (Luo and Kharas, 2024; Marcel et al., 2013), while its downregulation is also linked to adverse outcomes in breast cancer (Nguyen Van Long et al., 2022). Similarly, dyskerin overexpression has been observed in breast and prostate cancers (Kan et al., 2021; Stockert et al., 2019), whereas reduced levels are associated with breast and endometrial cancer progression (Alnafakh et al., 2021; Zacchini et al., 2022).

The essential role of CEBPZ, NOC2L, and NOC3L genes in vertebrate development (e.g., embryonic lethality in knockout mice) and hematopoiesis (as shown in zebrafish) and the control of genes of RNA metabolism, also highlights their importance beyond basic ribosome assembly, potentially linking them to pathways that control RNA-DNA hybrid resolution or modification. Using a cross-species approach, we show these genes are crucial for rRNA maturation and overall ribosomal activity. Notably, our findings indicate that NOC family members do not functionally compensate for one another, suggesting that each protein contributes uniquely to RNA processing. This non-redundant function is critical for cellular growth and viability in *Drosophila* (Destefanis et al., 2022) and human cells, emphasizing their fundamental and conserved role in maintaining ribosomal homeostasis. Indeed, we observed that a significant proportion (∼20%) of genes correlating with CEBPZ and NOC2L expression encode for regulators or participants in R-loop metabolism (Table 1, highlighted in red). In support of this, recent proteomic studies have identified CEBPZ as a component of protein complexes associated with R-loops (Wu et al., 2021), RNA–DNA hybrid structures that typically form during transcription. If not properly resolved, R-loops can induce RNA stress and DNA damage, contributing to genome instability (Petermann, Lan and Zou, 2022; Wu et al., 2021). This connection may imply context-dependent roles, where elevated expression of CEBPZ could represent a compensatory response to stress or ribosomal imbalance rather than direct oncogenic activity. However, its downregulation may indirectly promote genomic instability, a hallmark of cancer. Notably, R-loops can also form within the nucleolus due to the intense transcriptional activity of rRNA genes (Feng and Manley, 2022; Petermann, Lan and Zou, 2022; Wells, White and Stirling, 2019).

In conclusion, we have identified CEBPZ, NOC2L, and NOC3L as novel regulators of rRNA processing and ribosome biogenesis. Their expression levels in tumors vary significantly depending on cellular and tissue context. High expression is observed in most tumor types and is typically associated with aggressive tumor behavior. In contrast, low expression in renal cancers, mainly KICH and AML, correlates with advanced tumor stages. These findings suggest that rRNA dysregulation influences cancer progression through distinct context-dependent mechanisms shaped by the cellular environment and tissue type. Assessing the expression of these genes in physiological tissues and specific tumor contexts could provide valuable insights into their role in ribosomal biogenesis in cancer progression and aid in the development of targeted diagnostic and prognostic biomarkers.

## Materials and Methods

### Fly husbandry

Fly cultures and crosses were raised at 25 °C on a standard medium containing 9 g/L agar (ZN5 B and V), 75 g/L corn flour, 60 g/L white sugar, 30 g/L brewers’ yeast (Fisher Scientific), 50 g/L fresh yeast, 50 mL/L molasses (Naturitas), along with nipagin, and propionic acid (Fisher). The lines used, *NOC1-RNAi* (B25992), *NOC2-RNAi* (B50907), *and NOC3-RNAi* (B61872), were obtained from the Bloomington Drosophila Stock Center.

### Western blot and SUnSET assay

At 48 hrs after transfection with the relative siRNAs, cells were incubated with medium containing 10 % serum and puromycin at 1 µg/ml (Invitrogen, Thermo Fisher Scientific) for 40 min at room temperature, then recovered in 10 % serum/medium without puromycin for 30 min at room temperature. Cells were lysed in RIPA buffer supplemented with protease and phosphatase inhibitors. Protein concentrations were determined using the BCA assay (Thermo Fisher Scientific). Equal amounts of protein (40 µg) were separated by 10% SDS-PAGE and blotted using anti-puromycin antibodies (Sigma-Aldrich). Total protein loading was analyzed using stain-free technologies (TGX Stain-Free Fastcast). Signal detection was performed using an ECL substrate (GE Healthcare) and visualized using a ChemiDoc imaging system (Bio-Rad). Other Antibodies used were anti-p53 (Novocastra), RPL14, and mouse anti-beta-actin (Sigma Aldrich).

### Cell culture, RNAi treatment

HEK-293FT and HepG2 cells (ThermoFisher Scientific) were cultured in DMEM (Corning Inc.) supplemented with 10 % FBS, 2 mM L-glutamine, 100 U/ml penicillin, and 100 μg/ml streptomycin (all from Sigma-Aldrich) and maintained at 37 °C, 5 % CO_2_ in a humidified incubator. For RNA interference, cells were transfected with Lipofectamine RNAiMAX (ThermoFisher Scientific) following the manufacturer’s specifications. Transfected siRNA sequences were as follows: siRNA CEBPZ 1: rCrArArArArGrUrCrArGrUrArCrUrArArArArArArArGrCAA; siRNA CEBPZ 2: rUrUrGrCrUrUrUrUrUrUrArGrUrArCrUrGrArCrUrUrGrArG; negative control: scrambled negative control DsiRNA all from Integrated DNA Technologies. 72 hours after transfection, the cells were harvested, and total RNA was extracted.

### Cell growth and the clonogenic assay

HepG2 cells were plated in six-well plates after 48 hrs of transfection at the same concentration of 800 cells/ml and 400 cells/ml, for the siSCR and siCEBPZ in duplicates. Cells were fixed for and stained on days 1, 3, and 5 with a methanol/trypan blue solution (Sigma-Aldrich) for 30 minutes, then washed with water to remove excess staining. Photos were taken using a Zeiss Axio Imager M2 microscope. Four different frames/areas from each well were taken, and cells were counted from photos using ImageJ and data graphed using GraphPad Prism v10. Clonogenic efficiency was calculated based on the initial number of cells plated at day 0, then counted at days 1,3, and 5 (Misra and Rajawat, 2021).

### RNA extraction and RT-PCR

RNA was extracted from *Drosophila* larvae or HEK 293FT and HepG2 human cancer cells using the RNeasy Mini Kit (Qiagen) following the manufacturer’s instructions. The isolated RNA was quantified using a Nanodrop2000. Total RNA (1000 ng of total RNA was reverse-transcribed into complementary DNA (cDNA) using SuperScript IV VILO Master Mix (Invitrogen). The obtained cDNA was used for qRT-PCR with the SYBR Green PCR Kit (Qiagen). The assays were performed on a Bio-Rad CFX96 machine and analyzed using Bio-Rad CFX Manager software. Transcript abundance was normalized using *actin5c.* The primer list was published in (Destefanis et al., 2022). For human targets, primers were purchased from Integrated DNA Technologies: CEBPZ F: TCTCATCCAAAGTAGCCAGCAT, R: TCTCATCCAAAGTAGCCAGCAT; 45S pre-rRNA F: GAACGGTGGTGTGTCGTTC, R: GCGTCTCGTCTCGTCTCACT); p21 F: TGGGGATGTCCGTCAGAACC, R: TGGAGTGGTAGAAATCTCTCATGCT; BCL2 F: ATCGCCCTGTGGATGACTGAGT, R: GCCAGGAGAAATCAAACAGAGGC; BAX F: ATGTTTTCTGACGGCAACTTC, R: ATCAGTTCCGGCACCTTG, and GUSB housekeeping expression kit was purchased from Applied Biosystems (ref 4326320E).

### Statistical Analysis

Students’ *t*-test analysis and analysis of variance were calculated using one-way ANOVA, and Tukey’s multiple comparisons test was calculated using GraphPad-PRISM*8. P* values are indicated with asterisks ***\* =*** *P < 0.05, ** = P < 0.01, *** = P < 0.001, **** = P < 0.0001*, respectively.

### Correlations

TCGA data were accessed through FireBrowse (Broad Institute TCGA Genome Data Analysis Center, 2016): Firehose 2016-01-28 run. Broad Institute of MIT and Harvard. doi:10.7908/C11G0KM9) and processed in-house using the r-corr function of the Hmisc package in R. (https://www.rdocumentation.org/packages/Hmisc/versions/5.1-3) ^47^

### Differential expression and HR heat-map

Differential expression analysis was conducted using the GEPIA2 (Tang et al., 2019) platform to compare gene expression profiles between tumor and normal samples derived from the TCGA and GTEx datasets. GEPIA2 was also used to produce a survival heat map of hazard ratio.

### Gene Ontology enrichment

GO and KEGG enrichment analyses were performed using ShinyGO (Ge, Jung and Yao, 2020) with an FDR cutoff of 0.5. The shared coregulated genes between CEBPZ, NOC2L, and NOC3L were tested.

### Co-expressed genes

Broad DepMap https://depmap.org/portal/(Project data public release 24Q4) was used to determine the genes’ dependencies after CRISPR in cancer cell lines (Fong et al., 2024; Meyers et al., 2017; Tsherniak et al., 2017). The Broad DepMap project reports essentiality scores using the Chronos algorithm (Dempster et al., 2021). A lower score indicates a greater probability that the gene of interest is essential in a specific cell line. A score of 0 denotes a non-essential gene, whereas a score of −1 reflects the median for pan-essential genes.

### Tumor stage expression

The expression levels of the three genes of interest have been explored through UALCAN at https://ualcan.path.uab.edu ^32, 33^

## Acknowledgments

We thank the Bloomington Stock Center (NIH P40OD018537). Department CIBIO Core Facilities is supported by the European Regional Development Fund (FESR) 2021– 2027. This article is based on work from COST Action CA21154 TRANSLACORE, supported by COST (European Cooperation in Science and Technology and by the Dipartimento di Eccellenza 2023-2027, Legge 232/2016 project n 40613, funded by the MUR.

## Author contributions

G.R. performed the bioinformatics analysis. V.M. performed the Drosophila experiments. A.VC. analyzed protein homology and performed structural modeling, molecular simulations, and AlphaFolding. M.P. performed the analysis of CEBPZ expression and contributed to the final version of the manuscript. G.B. analyzed and contributed to the drafting of the data expression in tumors. P.B. drafted the initial manuscript, conceived the study, participated in its design, coordinated and supervised the team, and edited the final manuscript. All authors have read and agreed to the published version of the manuscript.

## Data availability statement

Data supporting the study for Figure 2 and Supplementary File 3 are available in DepMap (Tsherniak et al., 2017) at the URL https://depmap.org/portal using the keywords: CEBPZ, https://depmap.org/portal/gene/CEBPZ?tab=overview Top 100 co-dependencies.NOC2L, https://depmap.org/portal/gene/NOC2L?tab=overview Top 100 co-dependencies.NOC3L, https://depmap.org/portal/gene/NOC3L?tab=overview Top 100 co-dependencies. DepMap, Broad (2024). DepMap 24Q4 Public. Figshare+. Dataset. https://doi.org/10.25452/figshare.plus.24667905.v2 (Fong et al., 2024; Tsherniak et al., 2017).

Data supporting the study in Figures 3 and 4 are available from UALCAN (Chandrashekar et al., 2017; Chandrashekar et al., 2022) at https://ualcan.path.uab.edu/openly. Using the keywords: CEBPZ, NOC2L, and NOC3L at https://ualcan.path.uab.edu/cgi-bin/ualcan-res.pl. Then, select the respective expression from the tumors in the list. Gepia - GEPIA2 at http://gepia2.cancer-pku.cn/#index using CEBPZ, NOC2L, and NOC3L.

