## Supplementary material for "From Flies to Humans: A Conserved Role of CEBPZ, NOC2L, and NOC3L in rRNA Processing and Tumorigenesis": Supplemenraty file

### Running title

Conserved Roles of NOC Proteins in rRNA Processing and Tumorigenesis

### Keywords

NOC1/2/3, CEBPZ, NOC2L/3L, Ribosome biogenesis, rRNA processing, Nucleolar proteins, Cancer, Evolutionary conservation, *Drosophila*.

(Table 1 and in Figure 1P). In further maturation steps, CEBPZ detaches from NOC2L, allowing NOC3L to bind and heterodimerize with NOC2L in a complex necessary at state C of the 60S ribosomal maturation (Hurt, Iwasa and Beckmann, 2024). In addition to FTSJ3 and PES1, this complex contains NIP7 and DIS3, which were found to be coregulated with NOC2L-NOC3L in our expression analysis (Table 1 and in Figure 1P).

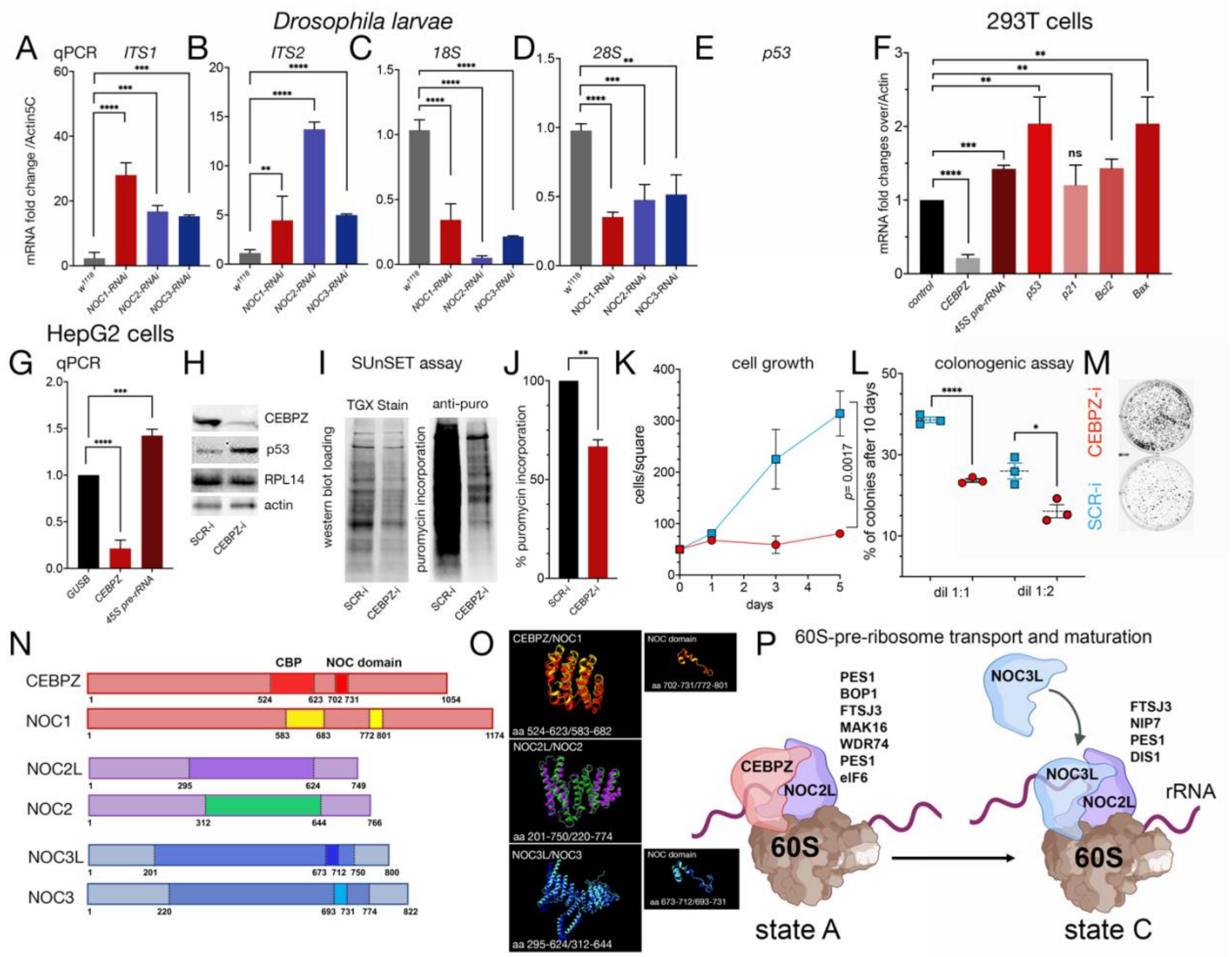


**Figure 1. (A-F) Reduction of Nucleolar Complex proteins (NOCs) in Drosophila leads to rRNA accumulation, impaired maturation of 18S and 28S rRNA, and elevated p53 levels**. (A-E) qRT-PCRs from *Drosophila* whole larvae with ubiquitous reduced *NOC1, NOC2,* or *NOC3* showing an accumulation of pre-rRNAs, analyzed using the ITS1 (internal transcribed spacers) 1 and 2 (A), and a reduction of 18S and 28S rRNA expression (C-D) and of *p53 mRNA* (E). The level of *NOC1, NOC2,* or *NOC3-mRNAs* is calculated over control $w^{1118}$ animals; data are expressed as fold increase relative to control *Actin5C;* RNA interference efficiency is shown in Supplementary Figure 1. **(F-M) Reducing CEBPZ in human tumor cells results in rRNA accumulation, impaired protein synthesis, and decreased cell growth.** (F-G) qRT-PCR analysis of HEK 293FT cells (F) and HepG2 cells (G) following siRNA-mediated silencing of CEBPZ, showing increased levels of 45S pre-rRNA and p53 mRNA. Expression levels are presented as fold change relative to control, normalized to β-glucuronidase (GUSB). Data from (A-G) are presented as mean ± s.d. for at least three independent experiments ***p*<0.01; ****p*<0.001; *****p*<0.0001 (using Student's t-test for the analysis). (H) Western blot from HepG2 lysates of cells transfected with a scramble siRNA (SCR-i) or for siCEBPZ (CEBPZ-i) showing the level of CEBPZ, p53, RPL14 used as an unrelated protein, and actin as a control loading. (I) SUnSET assay; western blot from cells treated

with 1 µg/ml of puromycin shows the relative changes in protein synthesis using anti-puromycin antibodies in cells treated with siSCR, or siCEBPZ, total protein loading is shown using stain-free technologies (TGX Stain-Free Fastcast). (J) Quantification of the relative change in the rate of puromycin incorporation between SCR-i and CEBPZ-i data was analyzed using two independent experiments. (K-L) (K) Cell growth expressed at day 1,3 and 5, and (L) percentage of colonies at 10 days after treatment with siCEBPZ and siCTL data are expressed from two separate experiments where cells were originally plated at two different concentrations *$p<0.05$; ****$p<0.0001$ (using Student's t-test for the analysis). (M) Photos of colonies at 10 days of treatment. **(N-O) Schematic representation of *Drosophila* and humans NOC1/CEBPZ, NOC2/NOC2L, and NOC3/NOC3L proteins.** CEBPZ contains a CBP (CCAAT binding domain in dark RED) with 32% amino acid-sequence identity with *Drosophila* NOC1, and a conserved NOC domain (in red and yellow). This domain is also present in NOC3L and NOC3 (highlighted in blue and azure). NOC2 shares an overall 36% of amino acid identity between *Drosophila* and human NOC2L, with the highest homology (47%) represented with a box (in purple and green); see sequence homologies in Supplementary File 2. **(O) The predicted structural homology of the conserved regions between CEBPZ/NOC1, NOC2L/NOC2, and NOC3L/NOC3** was obtained by a simulation using AlphaFold and ChimeraX analysis. **(P) Schematic representation of the CEBPZ-NOC2L and NOC2L-NOC3L heterodimers at states A and C of the 60S ribosomal maturation**. The graph also includes proteins identified through DepMap analysis (Table 1) due to their significant correlation with the expression of the corresponding heterodimers.

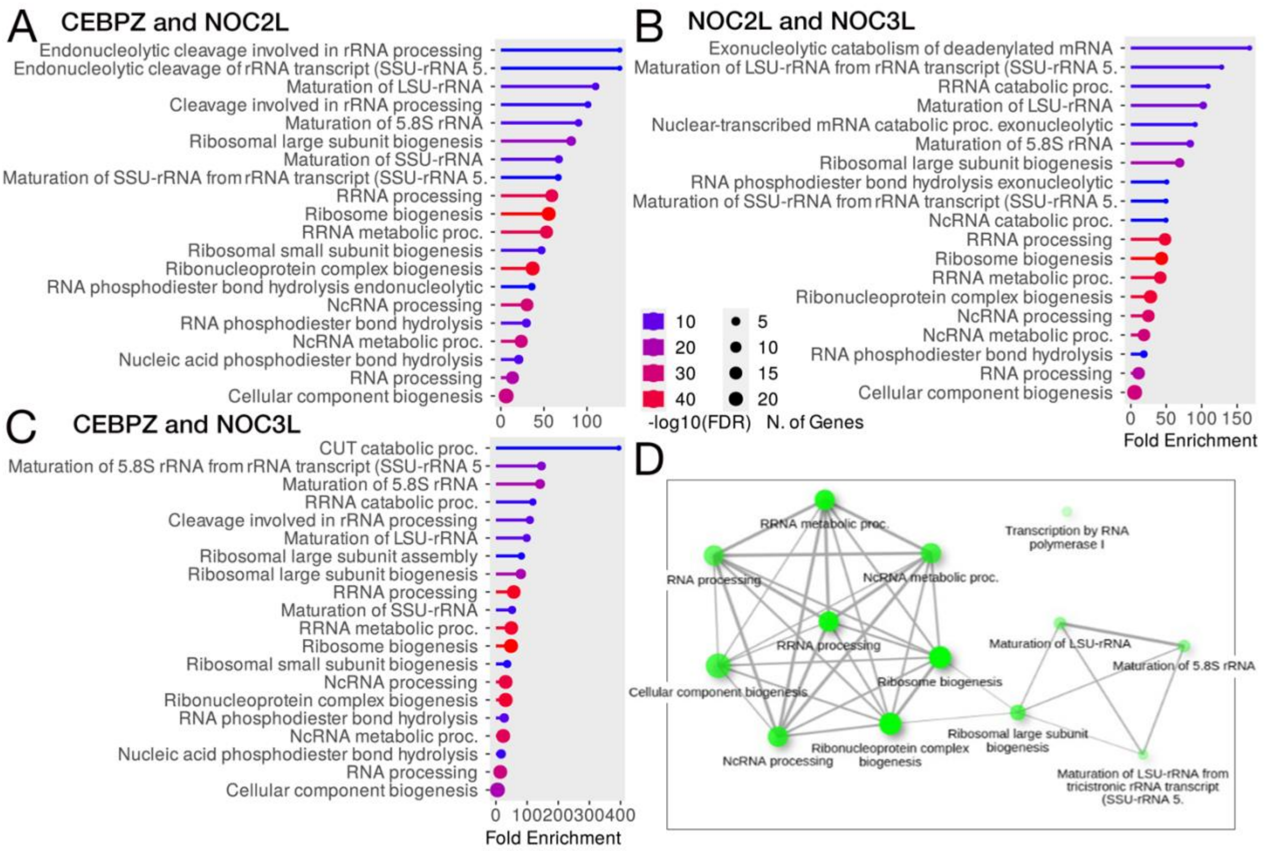


**Figure 2. GO enrichment of the shared co-regulated genes between CEBPZ, NOC2L, and NOC3L.** (A-C) GO enrichment (Biological Processes) of the shared co-regulated genes between the two indicated target genes. The results are ordered based on the Fold Enrichment; the circle size represents the number of genes in the pathway, the color of the bars the -log10(FRD), and the length of the bar the enrichment. (D) GO network representation. Brighter nodes are more significantly enriched gene sets. Bigger nodes represent larger gene sets. Thicker edges represent a higher number of genes shared between the two sets.

In Table 1, we show the list of genes that positively correlate with CEBPZ and NOC2L reduction, and we highlight in red the genes associated with R-loop formation of metabolism (22% of the total), excluding CEBPZ.

| Name | Function of the genes positively correlated with CEBPZ and NOC2L reduction |
|---|---|
| BOP1 | Component of the PeBoW complex, required for maturation of 28S and 5.8S ribosomal RNAs. |
| **DDX21** | RNA helicase acts as a sensor of the transcriptional status of RNA polymerase (Pol) I and II: promotes ribosomal RNA (rRNA) processing and transcription from polymerase II (Pol II) |
| DDX51 | ATP-binding RNA helicase involved in the biogenesis of 60S subunits. |
| **DDX56*** | Nucleolar RNA helicase that controls nucleolar integrity and RiBi |
| DHX33 | Implicated in nucleolar organization, stimulates RNA polymerase I transcription of the 47S precursor rRNA. Associates with ribosomal DNA (rDNA) loci where it is involved in POLR1A recruitment |
| DIMT1 | Demethylates two adjacent adenosines in the loop of a conserved hairpin near the 3'-end of 18S rRNA in the 40S particle. Involved in the pre-rRNA processing leading to small subunit rRNA. |
| DIS3** | Putative catalytic component of the RNA exosome complex 3'->5' exoribonuclease activity. |
| DKC1 | small nucleolar ribonucleoprotein (H/ACA snoRNP) complex, catalyzes pseudouridylation of rRNA. |
| EIF6 | Binds to the 60S subunit and prevents its association with the 40S to form the 80S |
| FTSJ3 | RNA 2'-O-methyltransferase involved in the 34S pre-rRNA to 18S rRNA and in 40S formation. |
| GRWD1 | Histone binding-protein that regulates chromatin dynamics and mini chromosome maintenance. |
| IMP4 | Component of the 60-80S U3 small nucleolar ribonucleoprotein (U3 snoRNP). |
| ISG20L2 | 3'-> 5'-exoribonuclease involved in ribosome biogenesis in the processing of the 12S pre-rRNA. |
| LAS1L | Required for the synthesis of the 60S ribosomal subunit and maturation of the 28S rRNA. |
| MAK16 | Important for the maturation of LSU-rRNA and 5.8S rRNA |
| MYBBP1A | May activate or repress transcription via interactions with sequence specific DNA-binding proteins. |
| NAT10 | RNA cytidine acetyltransferase catalyzes $N_4$-acetylcytidine modification on18S and rRNA. |
| NHP2 | Required for ribosome biogenesis and telomere maintenance. Part of the H/ACA small nucleolar ribonucleoprotein complex, which catalyzes pseudouridylation of rRNA. |
| **NIP7** | Required for 34S pre-rRNA processing and 60S ribosome assembly. |
| NOC4L | Nucleolar complex-associated protein 4-like protein. |
| **NOL12*** | RNA binding protein that plays a role in RNA metabolism, the resolution of DNA stress, nucleolar organization, regulates the levels of nucleolar fibrillarin and nucleolin in pre-rRNA processing. |
| NOL9 | Involved in rRNA processing, for the processing of the 32S precursor into 5.8S and 28S rRNAs. |
| NOP16** | Involved in the biogenesis of the 60S ribosomal subunit. |
| PAK1IP1 | Negatively regulates the PAK1 kinase. |
| PDCD11 | Essential for the generation of mature 18S rRNA, for cleavages at sites A0, 1 and 2 of the 47S. |
| PELP1* | Component of the PELP1 complex involved in the 28S rRNA maturation and transit of the pre-60S. |
| **PES1**** | PeBoW complex, required for the maturation of 28S and 5.8S RNAs and formation of the 60S. |
| POLR1E | Component of RNA polymerase I polymerase which synthesizes ribosomal RNA precursor. |
| POLR1G | Component of RNA polymerase I (Pol I) which synthesizes ribosomal RNA precursors. |
| **POP5** | Component of ribonuclease P, that generates mature tRNA molecules by cleaving their 5'-ends. |
| PPAN | A chimeric transcript, characterized by the first third of PPAN exon 12 joined to P2RY11 exon 2. |
| PWP1 | Regulates Pol I-mediated rRNA biogenesis and, probably, Pol III-mediated transcription. |
| PWP2 | Part of the small subunit (SSU) processome, precursor of the small eukaryotic ribosomal subunit. |
| RPL13A | Associated with ribosomes but is not required for canonical ribosome function. |
| RPP14* | ribonucleoprotein complex that generates mature tRNA molecules. |
| **RPP38** | Component of ribonuclease P complex, generates mature tRNA molecules by cleaving their 5'-ends. |
| RRP1* | Critical role in the generation of 28S rRNA |
| RRP12 | Required for nuclear export of both pre-40S and pre-60S subunits. |
| **TAF1C** | Component of the transcription factor SL1/TIF-IB complex, which is involved in the assembly of the PIC (pre-initiation complex) during RNA polymerase I-dependent transcription. |
| TBL3 | Part of the small subunit (SSU) processome, precursor of the small eukaryotic ribosomal subunit. |
| TIMM50** | Component of the TIM23 complex, mediates the translocation of transit peptide-containing proteins. |

| | |
|---|---|
| URB2 | Essential for hematopoietic stem cell development through the regulation of p53/TP53 pathway. |
| UTP20 | Part of the small subunit (SSU) processome, precursor of the small eukaryotic ribosomal subunit. |
| UTP23 | Involved in rRNA-processing and ribosome biogenesis. |
| UTP4 | Nucleolar processing of pre-18S ribosomal RNA. Part of the small subunit (SSU) processome. |
| WDR18 | Component of the PELP1 complex involved in the 28S rRNA maturation and transit of the pre-60S. |
| **WDR36** | Part of the small subunit (SSU) precursor of the small eukaryotic ribosomal subunit. |
| **WDR74** | Regulatory protein of the MTREX-exosome complex involved in the synthesis of the 60S rib.subunit. |
| WDR75 | Part of the small subunit (SSU) processome, precursor of the small eukaryotic ribosomal subunit. |

**Table 1. Genes positively correlated with CEBPZ and NOC2L upon their reduction in tumor cells and their functional annotation** from the UniProt database (UniProt, 2024).
* Positively correlating when CEBPZ and NOC3 are reduced; ** positively correlate when all three genes are reduced. The whole list of genes is available in Supplementary File 3. In red are genes associated with R-loop formation and metabolism.

### 3. CEBPZ, NOC2L and NOC3L tumor expression analysis.

Dysregulation of rRNA processing is a hallmark of cancers. We analyzed the expression across diverse tumor types based on the hypothesis that CEBPZ, NOC2L, and NOC3L form regulatory heterodimers that modulate this process. Specifically, we examined their expression levels using TCGA data from the UALCAN cancer portal (Chandrashekar et al., 2017; Chandrashekar et al., 2022) (Figure 3A-C). This analysis revealed that while all three genes were significantly upregulated ($P<0.05$) in most tumors (Figure 3D, in red), their expression was significantly reduced ($P<0.05$) in acute myeloid leukemia (LAML), and Kidney Chromophobe carcinoma (KICH) (Figure 3D, in green). In contrast, in KIRC and KIRP renal carcinomas, while the expression of CEBPZ was low, that of NOC2L and NOC3L increased (Figure 3D). These data provide evidence of different controls for the expression of these genes in different cell-tissue contexts. Indeed, as these genes may be upregulated in many solid tumors, to support increased ribosome production and protein synthesis for rapid cellular proliferation, their selective downregulation in hematopoiesis and kidney tumors KICH may suggest a tissue-specific or cell-specific regulation. Moreover, the fact that NOC2L and NOC3L are upregulated in KIRC/KIRP may indicate that they may exert a dual function, perhaps in transcriptional regulation or cell cycle control, beyond rRNA processing.

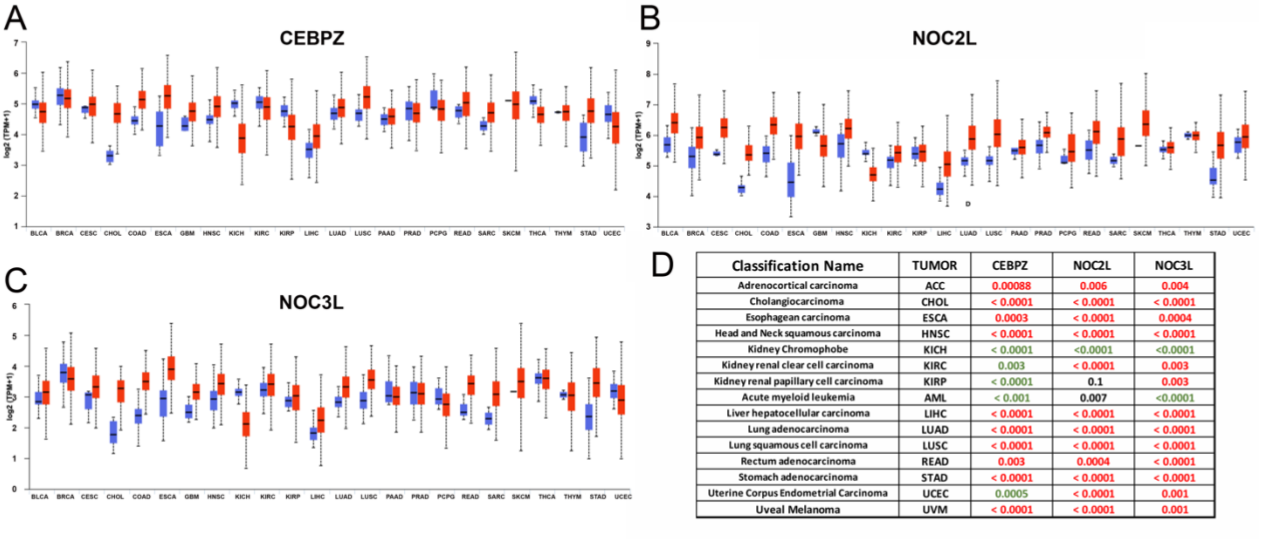


| Classification Name | TUMOR | CEBPZ | NOC2L | NOC3L |
|---|---|---|---|---|
| Adrenocortical carcinoma | ACC | 0.00088 | 0.006 | 0.004 |
| Cholangiocarcinoma | CHOL | < 0.0001 | < 0.0001 | < 0.0001 |
| Esophagean carcinoma | ESCA | 0.0003 | < 0.0001 | 0.0004 |
| Head and Neck squamous carcinoma | HNSC | < 0.0001 | < 0.0001 | < 0.0001 |
| Kidney Chromophobe | KICH | < 0.0001 | <0.0001 | <0.0001 |
| Kidney renal clear cell carcinoma | KIRC | 0.003 | < 0.0001 | 0.003 |
| Kidney renal papillary cell carcinoma | KIRP | < 0.0001 | 0.1 | 0.003 |
| Acute myeloid leukemia | AML | < 0.001 | 0.007 | <0.0001 |
| Liver hepatocellular carcinoma | LIHC | < 0.0001 | < 0.0001 | < 0.0001 |
| Lung adenocarcinoma | LUAD | < 0.0001 | < 0.0001 | < 0.0001 |
| Lung squamous cell carcinoma | LUSC | < 0.0001 | < 0.0001 | < 0.0001 |
| Rectum adenocarcinoma | READ | 0.003 | 0.0004 | < 0.0001 |
| Stomach adenocarcinoma | STAD | < 0.0001 | < 0.0001 | < 0.0001 |
| Uterine Corpus Endometrial Carcinoma | UCEC | 0.0005 | < 0.0001 | 0.001 |
| Uveal Melanoma | UVM | < 0.0001 | < 0.0001 | 0.001 |

**Figure 3. Gene expression analysis (TCGA).** (A-C) Expression levels of *CEBPZ* (A), *NOC2L* (B), and *NOC3L* (C) in tumor (red) versus normal (blue) tissues across various cancer types. (D) Summary panel highlighting cancer types where the expression of at least two of the three genes is significantly altered ($P < 0.05$), as reported in the UALCAN portal. Red indicates overexpression, while green indicates downregulation in tumor tissues relative to normal control.

We then examined CEBPZ*,* NOC2L*,* and NOC3L expression patterns across various stages of tumor development using TGCA datasets from the UALCAN portal. This analysis revealed a significant increase in their expression in liver hepatocellular carcinomas (LICH) and lung adenocarcinoma (LUAD) (Figure 4A-B). The observed upregulation correlated with the progression of tumor stages. The expression levels of LIHC in stage 4 were comparable to those in controls, possibly because of the reduced sample size for this stage. Similar upregulation was found for rectum adenocarcinoma (READ) and stomach adenocarcinoma (STAD) (not shown), suggesting that CEBPZ, NOC2L, and NOC3L overexpression may contribute to the aggressiveness of these carcinomas, probably by sustaining the high rate of rRNA processing and ribosome activity in these tumors. In contrast, in kidney chromophobe carcinomas, CEBPZ, NOC2L, and NOC3L expression levels were consistently and significantly reduced (Figure 4C), and variable in other kidney cancers (Figure 4D-E), supporting the conclusion of our previous analysis.

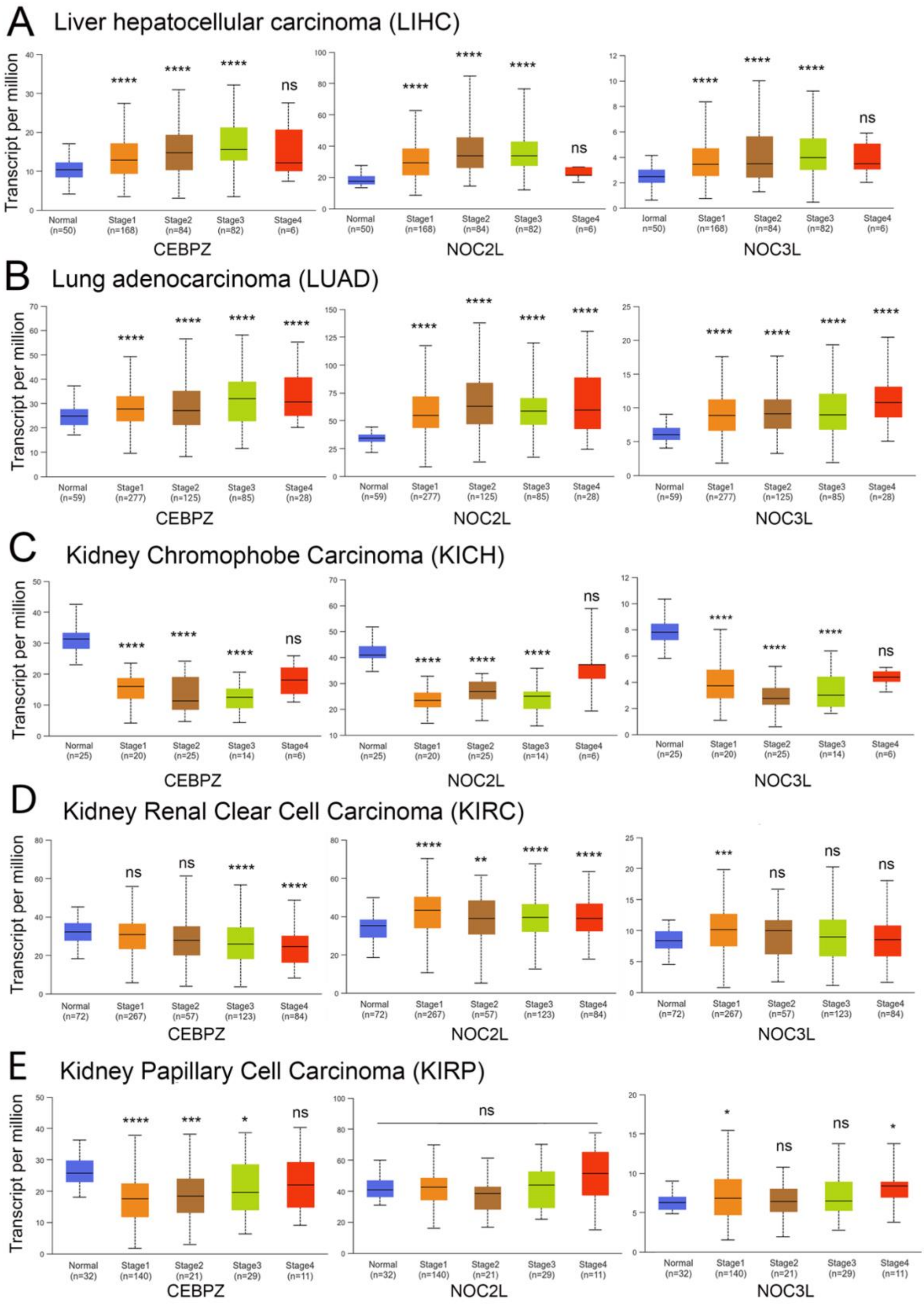


**Figure 4.** Graphs illustrating expression levels across various cancer types during tumor progression. Data from UALCAN highlights the correlation between gene expression and disease progression, demonstrating significant changes in the expression of CEBPZ, NOC2L, and NOC3L with advancing disease stages. (A) Liver hepatocellular carcinoma (HCC) and (B) Lung adenocarcinoma (LUAD), (C) Kidney Chromophobe Carcinoma (KICH). (D) Kidney Renal Clear

Cell Carcinoma (KIRC). (E) Kidney Papillary Cell Carcinoma (KIRP). The statistical analysis was performed by comparing the expression levels of the indicated genes in tumors at various stages to their expression levels in normal control tissue; statistical significance is expressed by the asterisks * $p< 0.05$, ** $p< 0.01$, *** $p< 0.001$, **** $p< 0.0001$, ns: not significant.

To deeply investigate the co-expression patterns of CEBPZ, NOC2L, and NOC3L within kidney tumors, we analyzed the TCGA datasets publicly accessible through FireBrowse (Cerami et al., 2012). A Pearson correlation coefficient was calculated between the expression of the three genes (CEBPZ, NOC2L, and NOC3L) in the three kidney carcinoma subtypes (KICH, KIRC, and KIRP). This analysis showed that CEBPZ and NOC3L exhibited the highest direct correlation in all three tumors (Table 2). Meanwhile, a weak negative correlation existed between CEBPZ and the other two genes in KIRC and KIRP. This suggests their potential distinct regulation across various kidney cancers.

| **KICH** | | | | **KIRC** | | | | **KIRP** | | | |
|---|---|---|---|---|---|---|---|---|---|---|---|
| r | | | | r | | | | r | | | |
| | CEBPZ | NOC2L | NOC3L | | CEBPZ | NOC2L | NOC3L | | CEBPZ | NOC2L | NOC3L |
| CEBPZ | 1 | -0,035481 | 0,7768214 | CEBPZ | 1 | -0,263372 | 0,5205663 | CEBPZ | 1 | -0,223414 | 0,4592963 |
| NOC2L | -0,035481 | 1 | 0,1116638 | NOC2L | -0,263372 | 1 | -0,225758 | NOC2L | -0,223414 | 1 | -0,151207 |
| NOC3L | 0,7768214 | 0,1116638 | 1 | NOC3L | 0,5205663 | -0,225758 | 1 | NOC3L | 0,4592963 | -0,151207 | 1 |
| *P* | | | | *P* | | | | *P* | | | |
| | CEBPZ | NOC2L | NOC3L | | CEBPZ | NOC2L | NOC3L | | CEBPZ | NOC2L | NOC3L |
| CEBPZ | NA | 0,7384625 | 0 | CEBPZ | NA | 4,50E-11 | 0 | CEBPZ | NA | 5,10E-05 | 0 |
| NOC2L | 0,7384625 | NA | 0,2919838 | NOC2L | 4,50E-11 | NA | 1,92E-08 | NOC2L | 5,10E-05 | NA | 0,0064759 |
| NOC3L | 0 | 0,2919838 | NA | NOC3L | 0 | 1,92E-08 | NA | NOC3L | 0 | 0,0064759 | NA |

**Table 2. A matrix highlighting the significant correlation between CEBPZ, NOC2L, and NOC3L expression in KICH, KIRC, and KIRP kidney carcinomas.** It shows that for each tumor type, the r (correlation coefficient) and the *P* value, highlighted in blue, are significant, with a correlation higher than -/+ 0.4. A higher absolute number means a stronger correlation between the samples' expression levels. (e. g. in KICH, when CEBPZ is low, NOC3L is also low). The lower part of the table shows the *P*-value for each correlation, highlighting the significant ones ($P<0.05$).

#### 3.2 Impact of CEBPZ, NOC2L, and NOC3L expression on patient survival.

Finally, we analyzed the correlation between the expression of CEBPZ, NOC2L, and NOC3L and patient survival to explore their potential impact on disease outcomes across tumor types. Figure 5 displays a heatmap of hazard ratios for these genes, providing a visual summary of how their expression is related to survival in various tumors.

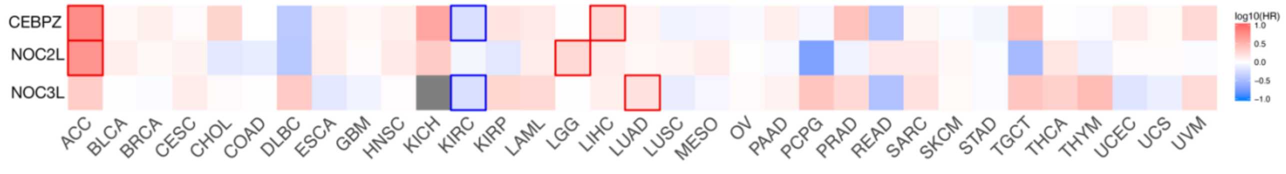


**Figure 5. Hazard ratio (HR) heat map for CEBPZ (ID: ENSG00000115816.13), NOC2L (ID: ENSG00000188976.10), and NOC3L (ID: ENSG00000173145.11) in the tumors listed below.** The median is selected as a threshold for separating high-expression and low-expression cohorts. The bounding boxes depict the statistically significant results ($P < 0.05$). Colors from white to red

indicate an HR above 1, suggesting that high gene expression is associated with a worse prognosis.

**Funding**

This work was supported by a NIH Public Health Service grant from NIH-SC1DK085047 to PB and Pallotti Legacy for Cancer Research to MP. GR is a recipient of a PhD fellowship in the PON program 'Research and Innovation' 2014-2020 with financial support from REACT-EU resources.
